# Multifunctional Materials Strategies for Enhanced Safety of Wireless, Skin-Interfaced Bioelectronic Devices

**DOI:** 10.1101/2023.02.28.530037

**Authors:** Claire Liu, Jin-Tae Kim, Da Som Yang, Donghwi Cho, Seonggwang Yoo, Surabhi R. Madhvapathy, Hyoyoung Jeong, Tianyu Yang, Haiwen Luan, Raudel Avila, Jihun Park, Yunyun Wu, Kennedy Bryant, Min Cho, JiYong Lee, Jay Kwak, WonHyoung Ryu, Yonggang Huang, Ralph G. Nuzzo, John A. Rogers

## Abstract

Many recently developed classes of wireless, skin-interfaced bioelectronic devices rely on conventional thermoset silicone elastomer materials, such as poly(dimethylsiloxane) (PDMS), as soft encapsulating structures around collections of electronic components, radio frequency antennas and, commonly, rechargeable batteries. In optimized layouts and device designs, these materials provide attractive features, most prominently in their gentle, noninvasive interfaces to the skin even at regions of high curvature and large natural deformations. Past work, however, overlooks opportunities for developing variants of these materials for multimodal means to enhance the safety of the devices against failure modes that range from mechanical damage to thermal runaway. This paper presents a self-healing PDMS dynamic covalent matrix embedded with chemistries that provide thermochromism, mechanochromism, strain-adaptive stiffening, and thermal insulation, as a collection of attributes relevant to safety. Demonstrations of this materials system and associated encapsulation strategy involve a wireless, skin-interfaced device that captures mechanoacoustic signatures of health status. The concepts introduced here can apply immediately to many other related bioelectronic devices.

## 1. Introduction

Wireless, skin-interfaced bioelectronic devices are increasingly popular for unobtrusive, noninvasive, and continuous monitoring of essential health parameters, ranging from traditional vital signs to various emerging metrics of patient status.^[1-5]^ The most mature of these devices consist of thin, flexible printed circuit boards (PCBs) structured into open mesh geometries with electronic components, sensing modules, and hardware for wireless communication. Power can be delivered using wireless schemes or provided by integrated batteries, which are encapsulated with the other constituent parts within a soft, flexible, and stretchable polymeric structure, typically of a medical-grade thermoset silicone elastomer. Ensuring that these wireless electronic device systems operate reliably and safely while adhered to the skin is a critical concern in engineering design. While many safeguards, spanning careful quality control to voltage/current protection circuits, are important, they do not address all possible safety hazards.^[3, 6-7]^ For example, mechanical failures in the encapsulation structure that follow from prolonged cycles of use can expose the user to mechanical and electrical hazards associated with the electronic components.^[8-12]^ Polymer chemistries that offer self-healing capabilities and strain-limiting features have advantages over simple silicones as encapsulants. Adding into these materials species for visual, colorimetric indication of these and other forms of device failure, are also of interest. As bioinspiration, the skins of chameleons provide many of the desired features: (1) self-healing capabilities, (2) colorimetric response to hazardous conditions, (3) variable mechanical stiffness and J-shaped stress/strain responses.^[13-17]^

Here, we report the development of materials with these attributes and modes for their use with wireless, skin-interfaced bioelectronic devices. Specifically, a self-healing poly(dimethylsiloxane) (SH PDMS) material serves as a foundational matrix for chemistries that support various stimuli-responsive functionalities and safety characteristics. Companion processing approaches provide routes for using these materials in soft, encapsulating structures as water-proof enclosures around multifunctional electronic systems. This strategy represents a materials-based complement to traditional safety mechanisms based on circuit designs and mechanical layouts. We illustrate the applicability of these schemes in miniaturized, wireless mechanoacoustic sensors, previously demonstrated for capturing a broad range of physiological parameters, along with bulk movements of the body.^[3-4]^ These results not only offer immediate, broad potential across wide ranging classes of devices, but also motivate the development of additional, complementary materials strategies for safe operation of bioelectronic systems.

## 2. Results and Discussion

### 2.1. Multifunctional Materials Design Approach

The operation of wireless, skin-interfaced bioelectronic devices involves intrinsic safety risks, such as those that may arise from mechanical damage to the encapsulating structure or an overheating battery (**Figure 1**A). The multifunctional materials design approach introduced here addresses these concerns through the use of a composite structural system that can reversibly and autonomically self-heal upon mechanical damage and change color upon certain excitative stimuli, such as temperature and strain. Additional features provide protection against excessive mechanical or thermal loads.

**Figure 1:**
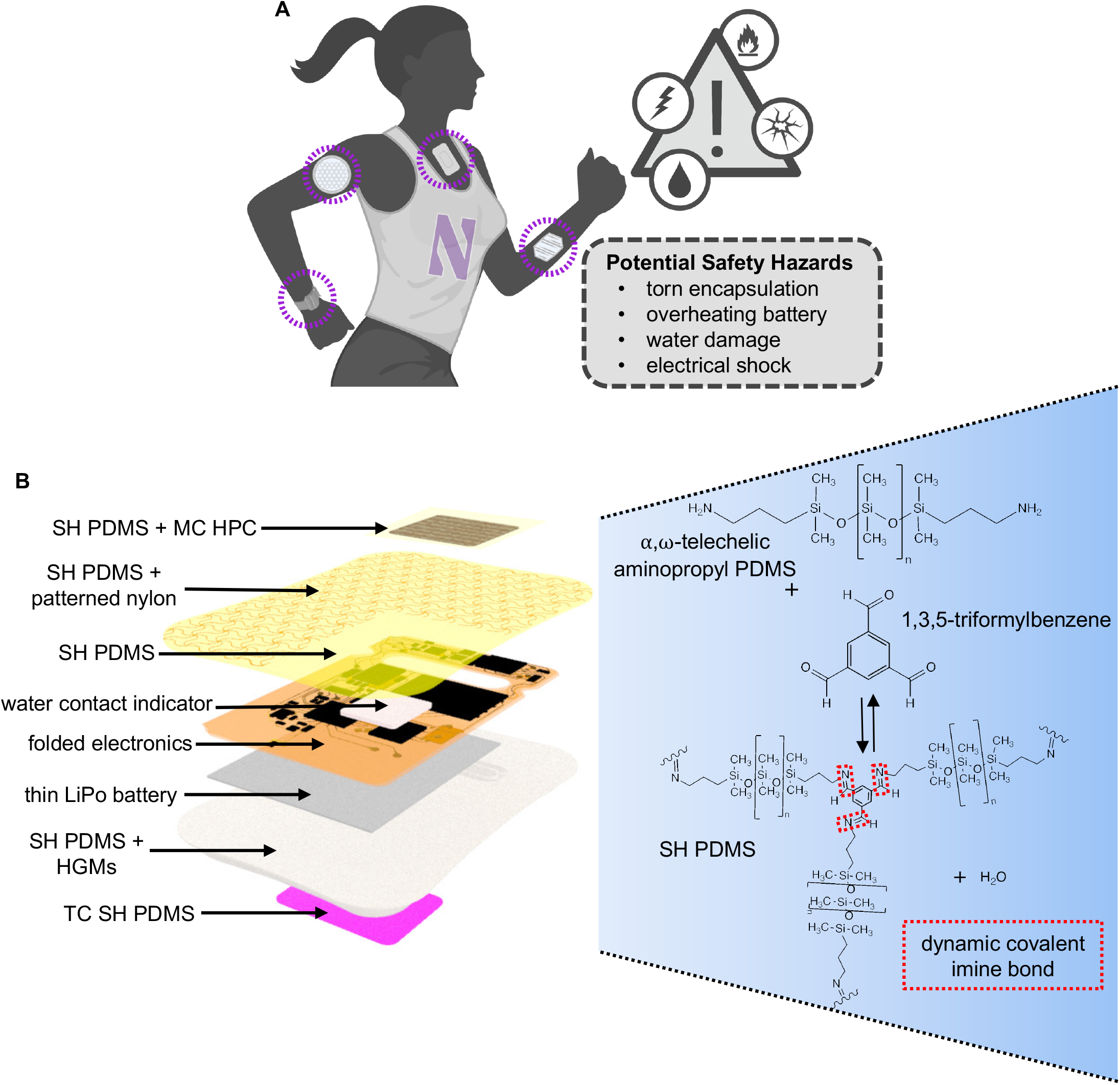
A multifunctional materials design approach for encapsulation of wireless, skin-interfaced bioelectronic devices with enhanced safety. **A)** Potential safety hazards associated with wireless, skin-interfaced bioelectronic devices. **B)** Schematic representation (left) of the structural configuration and the encapsulation material choices for the wireless, skin-interfaced bioelectronic device. All such encapsulation materials are based on self-healing poly(dimethylsiloxane) (SH PDMS), including thermochromic SH PDMS (TC SH PDMS), SH PDMS + hollow glass microspheres (HGMs), SH PDMS + mechanochromic hydroxypropyl cellulose (MC HPC), and SH PDMS + patterned nylon. The Schiff-base chemical reaction (right) between α,ω-telechelic aminopropyl PDMS and 1,3,5-triformylbenzene forms dynamic covalent imine bonds that serve as autonomic self-healing points in the SH PDMS elastomer matrix.

Self-healing properties within an elastomeric material, including commonly used silicones such as PDMS, can be achieved using extrinsic or intrinsic means.^[18]^ The former relies on capsules of healing agent distributed within an elastomer matrix, as reported by Sottos and Braun.^[19-20]^ Upon a damage event, the affected capsules break open to release a healing agent that flows into open cracks or other defects where it then solidifies. This method, while attractive, suffers from limited self-healing efficiency (i.e., the ability for the healed material to stretch to its original maximum elongation length) and supports only a single cycle of healing. The intrinsic form of self-healing relies on reversible, dynamic chemical bond formation within the material itself, thereby providing a reversible mechanism for healing.^[18, 21-22]^ Furthermore, given sufficient time for self-healing, the efficiency can be high.

These latter attractive features motivate the use of intrinsic self-healing concepts (Figure 1B)—in the form of a dynamic covalent SH PDMS formulation that features imine bonding as the primary mechanism for self-healing—as an advanced encapsulation material for skin-interfaced bioelectronic devices.^[23-25]^ Specifically, the chemistry relies on a one-pot, Schiff base condensation reaction between α,ω-telechelic aminopropyl PDMS and 1,3,5-triformylbenzene. We show that with additional chemical and structural components, including thermochromic (TC) leuco dyes, mechanochromic (MC) hydroxypropyl cellulose (HPC) sheets, patterned nylon meshes, and hollow glass microspheres (HGMs), it is possible to support a range of important stimuli-responsive properties within this SH PDMS matrix.

### 2.2. An Autonomic Self-Healing Dynamic Covalent Elastomer with Thermochromic (TC) and Mechanochromic (MC) Properties

The SH PDMS (**Figure 2**A-B) matrix described above consists of a light-yellow, transparent elastomer with dynamic covalent imine bonds for self-healing, as shown in characteristic peaks in the FTIR spectra (Figure 2C).^[23-25]^ Digital image correlation (DIC)^[26-27]^ experiments reveal that self-healing of SH PDMS occurs as soon as 15 minutes after a damage event (Figure S1). Self-healing can be quantified by the self-healing efficiency (η), defined as the ratio of the maximum strain of the self-healed sample to the maximum strain of the pristine sample (Table S1).^[23]^ After 15 minutes of self-healing from initial damage, while η_SH PDMS_ = 32.2%, the material still can sustain strain levels above 100%. Moreover, after 24 hours of self-healing, η_SH PDMS_ = 99.9% and the material is essentially restored to its pristine mechanical state.

**Figure 2:**
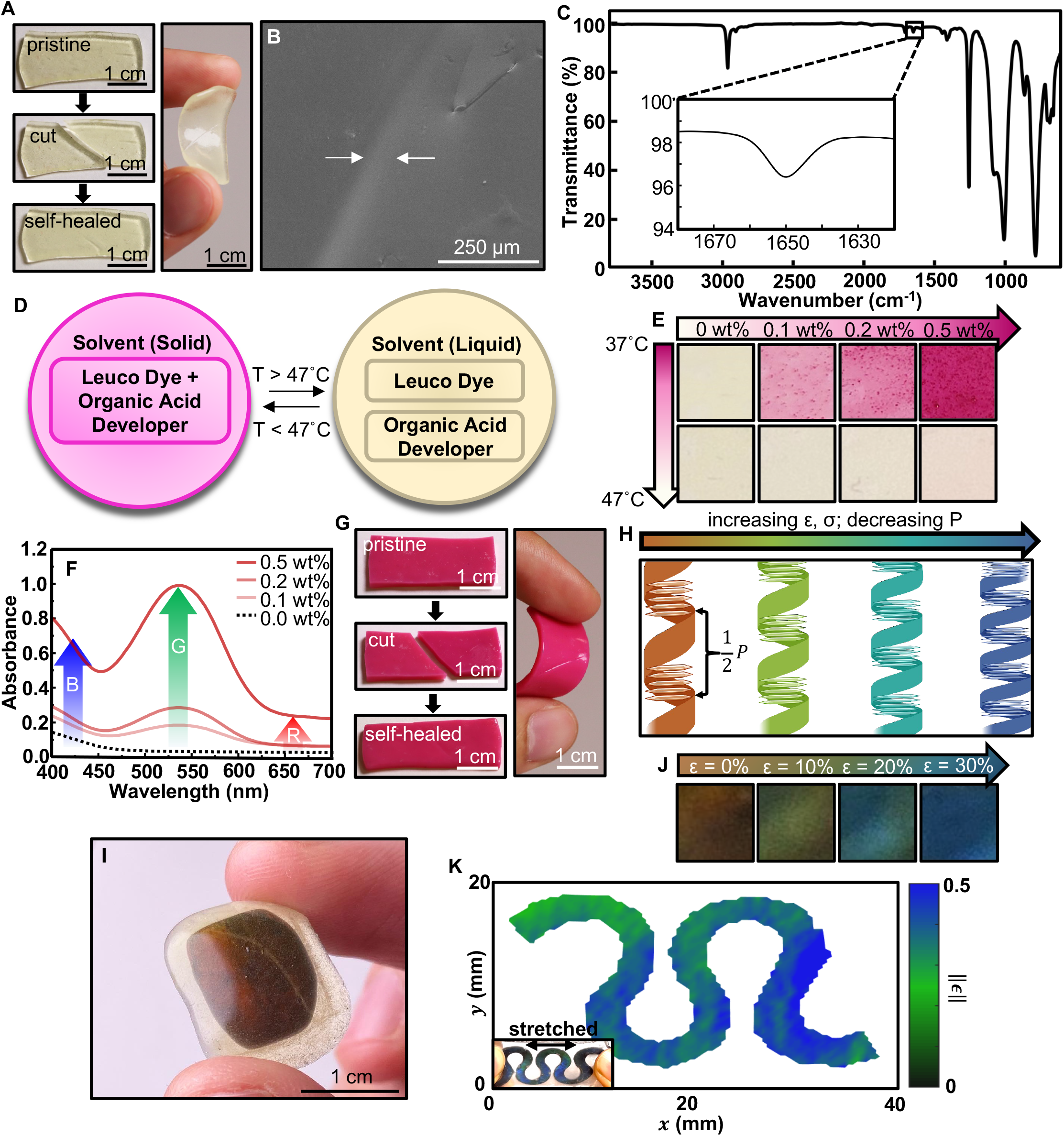
A dynamic covalent elastomer integrated with multiple stimuli-responsive and functional properties, including autonomic self-healing, thermochromism, and mechanochromism. **A)** Photographs illustrating the synthesized dynamic covalent elastomer (SH PDMS) self-healing after a damage event. **B)** A scanning electron microscopy image illustrating the self-healed region of the synthesized SH PDMS. **C)** FTIR spectra of the synthesized SH PDMS, illustrating the characteristic peak for the dynamic covalent imine bond. **D)** Schematic depiction of the thermochromic (TC) properties of a leuco dye (microcapsule form) with an activation temperature of 47°C, the temperature at which the dye switches from the colored to the colorless state. **E)** Color changes associated with a temperature increase from 37°C to 47°C of SH PDMS with varying mass fractions of the TC leuco dye. **F)** UV spectra of the SH PDMS with varying mass fractions of the TC leuco dye. **G)** Photographs illustrating the synthesized TC SH PDMS, integrated with 0.5 wt% TC leuco dye, undergoing autonomic self-healing after a damage event. **H)** Schematic depiction of the mechanochromic (MC) properties of the cholesteric liquid crystalline nanostructure of aqueous hydroxypropyl cellulose (HPC). **I)** Photograph illustrating the cholesteric liquid crystalline HPC encapsulated within thin, flexible layers of SH PDMS. **J)** Color changes associated with increasing mechanical strain of the encapsulated cholesteric liquid crystalline HPC. **K)** Digital image correlation experiments for simultaneous mechanical and chromic analysis of the encapsulated cholesteric liquid crystalline HPC, in a serpentine configuration, undergoing uniaxial strain.

Thermochromism represents an important safety feature in the context of device encapsulation, as a visual warning of overheating that can enable early removal of a device to minimize or eliminate collateral damage to the skin. Thermochromism in the SH PDMS matrix follows from incorporation of a microencapsulated leuco dye, tertiary component systems that consist of a dye (color former), developer, and solvent.^[28-29]^ The dye forms a colored complex with the developer at temperatures below the melting point of the solvent. At temperatures above this value, the phase change of the solvent destroys the colored complex between the dye and developer, thereby inducing a reversible change in color. The dye used here has a “switching temperature” of 47°C and a corresponding reversible change in color from magenta/pink to colorless (Figure 2D). This temperature represents the approximate value at which pain/discomfort is expected on human skin.^[6, 30-32]^ The International Electrotechnical Commission (IEC) defines 48°C as the maximum temperature that healthy adult human skin can be in contact with a medical device part, for contact times between 1-10 min.^[33]^ Various mass fractions of the leuco dye can be incorporated to yield desired properties (Figure 2E). Increasing the mass fraction of the dye to 0.5 wt% increases the visual vibrancy of the initial magenta state, as reflected in the increased absorbance for green wavelengths observed in the UV-Vis absorption spectra of the composite material (Figure 2F). This loading level also enables high color contrast (Figure S2). Therefore, a mass fraction of 0.5 wt% in the given film thickness (∼300 μm) provides the most effective thermochromism, as further loading generally degrades the uniformity of the composite material due to aggregation of the dye. This formulation, designated as TC SH PDMS, serves as the basis for device encapsulation results described subsequently. As shown in Figure 2G, the TC SH PDMS composite retains self-healing capabilities that are comparable to those of SH PDMS. After 24 hrs of self-healing, the TC SH PDMS composite achieves a healing efficiency of η_TC SH PDMS_ = 92.2% (Figure S3, Table S1).

Mechanochromism in SH PDMS can be realized using an aqueous solution of HPC, which enters the cholesteric liquid crystalline phase at a polymer weight percentage of at least 60%.^[34-36]^ The pitch size of the cholesteric liquid crystal helical nanostructure decreases as the material undergoes increasing strain/stress, thereby causing a blue-shift in the spectrum of the reflected light and a corresponding visible color change (Figure 2H). By integrating the MC HPC within thin layers of transparent SH PDMS (SH PDMS + MC HPC) (Figure 2I), the iridescent color changes as the material undergoes increasing levels of strain (Figure 2J). At strains of approximately ∼20-30%, the SH PDMS + MC HPC exhibits a significant blue-shift and the initial amber-hued color of the MC HPC component disappears, revealing vibrant green/blue tones (Figure 2J, Figure S4, Supplementary Video 1). DIC experiments quantify the displacement and strain distribution fields of a uniaxially stretched structure of this composite material in a serpentine shape (Figure 2K), enabling direct comparison with the mechanochromic behavior.^[26]^ The Triangular Cosserat Point Elements (TCPE) method reveals the strain distribution field, in a manner that allows separation of rigid body motions.^[27]^ These results (Figure 2K) are consistent with the color changes observed in the composite material (Figure 2K, inset), as further validated by finite element analysis (FEA) (Figures S5-S6).

### 2.3. An Autonomic, Self-Healing Dynamic Covalent Elastomer with Patterned Nylon Meshes for Strain-Adaptive Stiffening Properties

Skin, like many other biological tissues, exhibits an increase in stiffness with increasing strain, thereby reducing the potential for damage/injury under extreme applied forces.^[13]^ This unique mechanical property, characterized by the so-called “J-shaped” stress/strain response^[37-38]^, follows from its collagen/elastin composite structure and is not found in conventional silicone elastomers. The embedded elastin imparts elasticity, resulting in the initial linear stress/strain response and low modulus at low strain. At high strains, the collagen microfibers, which provide structural integrity and stiffness, begin to uncoil, straighten, and stretch. The result is a transition to a high-modulus regime and a corresponding increase in stiffness. Previous work reports the use of filamentary microstructures of polyimide embedded in low-modulus silicone elastomer gel to replicate the nonlinear mechanical stress/strain responses of biological tissues.^[39]^ The advances presented here extend this bioinspired materials design concept to materials for encapsulation. The approach uses high-throughput laser ablation methods to pattern triangular serpentine mesh structures in films of polyamide (Nylon-6, E_bulk_ ∼1.97 GPa, thickness ∼ 250 μm) that are then embedded in thin layers of SH PDMS as a composite structure (**Figure 3**A). The results are nonlinear stress/strain behaviors (Figure 3B). Corresponding photographs illustrate the structural deformations of a representative pattern as applied strain increases—transitioning from the original “toe” state, then to the “heel” state, and finally to the “linear” state. This scheme and design also apply to mesh structures of other types of polymers (Figure S7), including poly(ethylene terephthalate) (PET), poly(methyl methacrylate) (PMMA)), or even bioresorbable materials (e.g., cellulose acetate (CA), silk, poly-L-lactic acid (PLLA)).^[40-42]^

**Figure 3:**
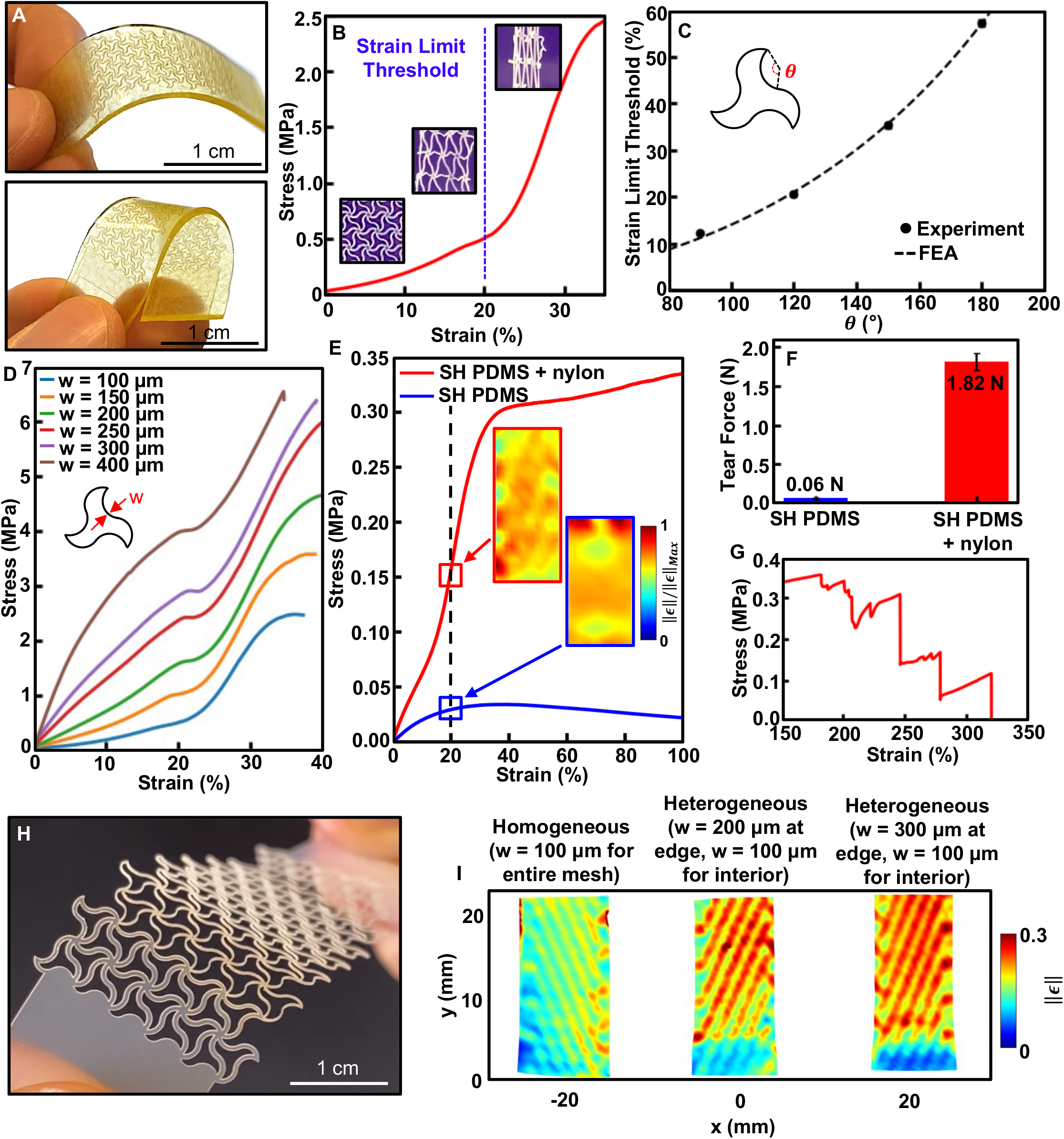
A dynamic covalent elastomer integrated with patterned nylon mesh mechanical reinforcement for nonlinear, strain-adaptive stiffening behavior. **A)** Photographs depicting the synthesized dynamic covalent elastomer (SH PDMS) integrated with a thin serpentine patterned nylon mesh for mechanical reinforcement. **B)** The nonlinear stress/strain profile of a thin serpentine patterned nylon mesh, along with corresponding photographs of the mesh. **C)** The strain limit threshold as a function of the nylon serpentine mesh angle (***θ***), with finite element analysis (FEA) modeling results. **D)** Stress/strain curves of the patterned nylon mesh (***θ*** = 120°) with varying serpentine widths (w). **E)** Stress/strain curve of the SH PDMS + patterned nylon mesh composite material, compared to SH PDMS without the patterned nylon mesh, along with corresponding strain distribution profiles obtained via digital image correlation (DIC) experiments. **F)** Tear force measurements of SH PDMS and SH PDMS + patterned nylon mesh composite material, as obtained from trouser tear-propagation testing. **G)** Stepwise tearing profile (stress/strain curve) of the SH PDMS + patterned nylon mesh composite material. **H)** Photograph of the heterogeneous serpentine patterned nylon mesh, featuring thicker serpentine widths at the edge. **I)** DIC experiments illustrate spatially varying strain distribution profiles of the SH PDMS + heterogeneous patterned nylon mesh composite material undergoing uniaxial strain.

For use on the skin, a strain limit threshold of ∼20%, corresponding to the strain at which the stiffness of the encapsulation material dramatically increases, is a reasonable design goal. This level of strain is also the point at which the SH PDMS + MC HPC composite material undergoes a strong blue-shift (Figures 2J-K). The serpentine mesh pattern determines the strain limit threshold response, via the serpentine angle (θ) (Figure 3C) and the serpentine width (w) (Figure 3D). For a given serpentine width (e.g., w = 100 μm), as θ increases from 90° to 180°, the strain limit threshold demonstrates an increasing trend, with θ = 120° yielding an average strain limit threshold of ∼20.5% (Figure 3C), a result further confirmed by FEA simulations. For a given serpentine angle (θ = 120°), as w increases from 100 to 400 μm (Figure 3D), the initial modulus at low levels of strain demonstrates an increasing trend, with the lowest modulus of E ∼ 1.08 MPa for w = 100 μm. This change represents a ∼96.7% and ∼99.9% reduction from that of w = 400 μm (E ∼ 33.02 MPa) and of bulk Nylon-6 (E ∼ 1.97 GPa), respectively. For a strain limit threshold of ∼20% and a low modulus response at strains below this threshold, therefore, the optimized parameters of the ∼250 μm-thick films are θ = 120° and w = 100 μm.

Figure 3E shows the nonlinear stress/strain behavior of the SH PDMS + nylon mesh composite material, as compared to SH PDMS. Indeed, the composite shows the desired strain-adaptive stiffening response, as the initial modulus of the composite material (E ∼ 0.82 MPa) increases by more than ∼90% to 1.58 MPa at ∼20% strain. Compared to SH PDMS (E ∼ 0.25 MPa), this mechanical reinforcement at ∼20% strain dramatically increases the modulus of the SH PDMS by ∼532%, to E ∼ 1.58 MPa.

The patterned mesh also increases the tear resistance, resulting in a ∼30x larger tear force (Figure 3F) and a stepwise tear profile (Figure 3G, Figure S8) that ultimately delays the complete fracture of the material. This behavior is consistent with conventional strategies that use randomly oriented fiber reinforcement networks.^[43-44]^ Moreover, the triangular geometry of the serpentine mesh lends itself to an inherent anisotropy that yields different strain threshold limits along different directions of strain (Figure S9). This unique property could be leveraged for applications that require differing levels of stretchability/mechanical reinforcement along the major axis vs. the minor axis of the device, such as a previously reported rectangular-shaped, wireless ECG device with electrodes that primarily stretch (∼20%) along the major axis.^[1-2]^

Another design option is to introduce spatial heterogeneity in the serpentine mesh design, such as increasing the serpentine width at the edge regions of the mesh (Figure 3H, Supplementary Video 2). Uniaxial stretching of such a mesh design encapsulated in SH PDMS yields a spatially varying strain distribution that illustrates lower strain at the edge regions—thereby endowing higher resistance to deformation at these regions (Figure 3I). Such design features may be relevant in the context of peeling/removal of the device from the skin, as there are typically greater stresses in these areas produced from handling of the device edges.

### 2.4. An Autonomic, Self-Healing Dynamic Covalent Elastomer with Hollow Glass Microspheres (HGMs) for Improved Thermophysical Properties

Thermally insulating properties are additional important aspects in safe encapsulating structures, to minimize heat transfer from overheating electronic components or batteries. HGMs, which consist of a thin outer borosilicate shell, are frequently utilized as inert additives in polymer matrices to reduce the thermal conductivity (*k*) and density (ρ) of the composite material.^[45-50]^ Small mass fractions (2.5 – 5.0 wt%) of HGMs integrated with SH PDMS yield a white appearance consistent with an expected increase in optical scattering (**Figure 4**A, Figure S10). Scanning electron microscopy (SEM) of fracture surfaces of composites with 2.5 and 5.0 wt% HGMs (Figure 4B) shows well-dispersed collections of HGMs within the SH PDMS matrix, in closed-shell morphologies. Uniaxial tensile testing (Figure 4C), via dynamic mechanical analysis (DMA), of SH PDMS + HGM composites reveals that SH PDMS and SH PDMS + HGM (2.5 wt%) have comparable levels of stretchability (>450% strain) and softness (E_SH PDMS_ ∼ 0.25 MPa; E_2.5wt% HGMs_ ∼ 0.32 MPa). For an HGM mass fraction of 5.0 wt%, the stretchability decreases significantly (<40% strain) and the stiffness of the composite material increases by ∼250% (E_5.0wt% HGMs_ ∼ 0.87 MPa) compared to SH PDMS. In both cases, these composites retain an ability to autonomically self-heal, comparable to SH PDMS. After a 24-hr period of autonomic self-healing, η_SH PDMS_ is 99.9%, η_2.5 wt% HGMs_ is 98.9%, and η_5.0 wt% HGMs_ is 82.0% (Table S1).

**Figure 4:**
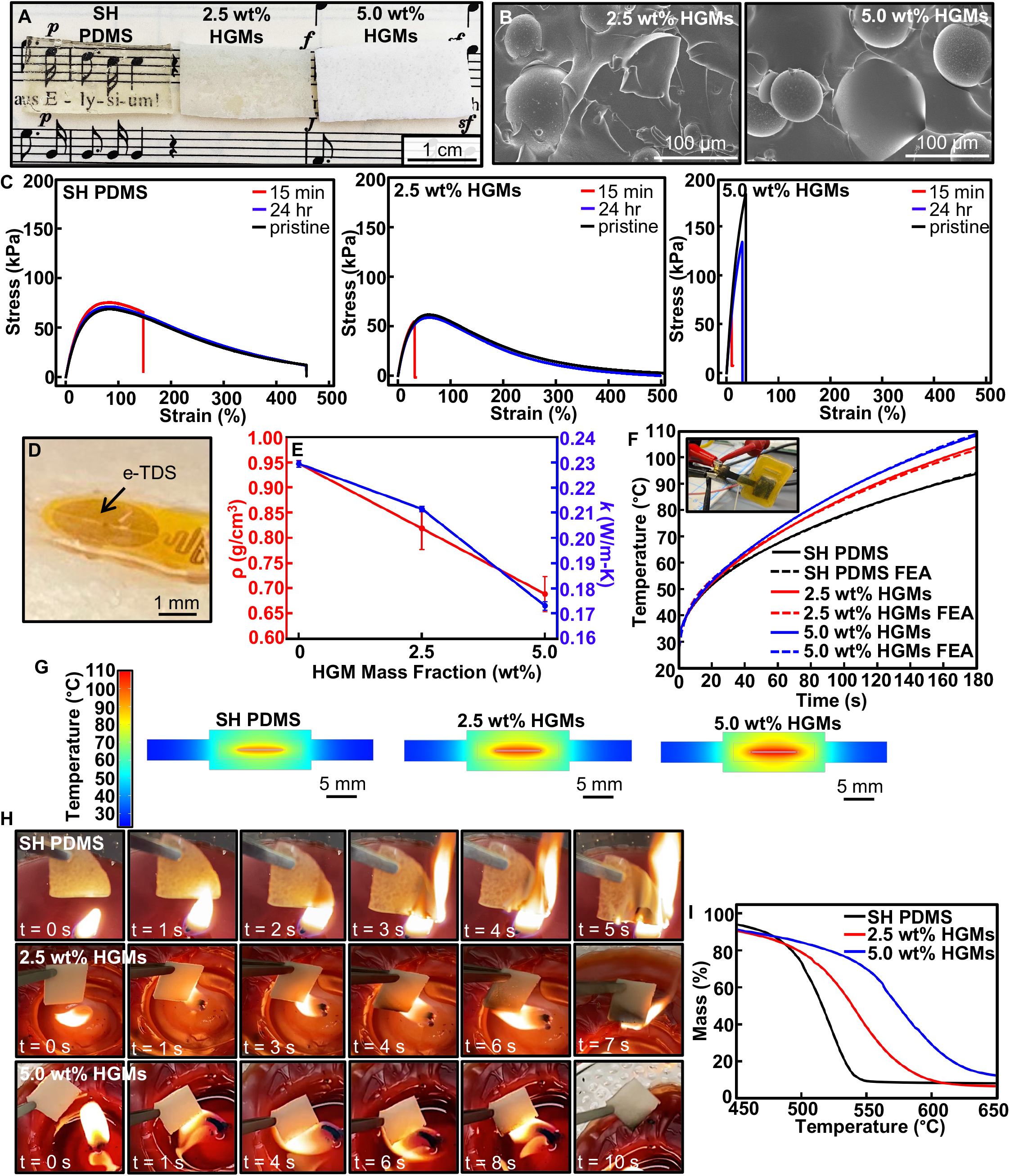
A dynamic covalent elastomer integrated with hollow glass microspheres (HGMs) to form an autonomic self-healing, low-density, and thermally insulating composite material. **A)** Photograph depicting the synthesized dynamic covalent elastomer (SH PDMS) integrated with varying mass fractions of HGMs. **B)** Scanning electron microscopy micrographs of the cross-sectional areas of SH PDMS + HGMs composites. **C)** Stress/strain curves of SH PDMS (left) and SH PDMS+ HGMs composites (middle and right), after various time periods for self-healing. **D)** Photograph illustrating an epidermal thermal depth sensor (e-TDS) on the top surface of the SH PDMS + HGMs composite. **E)** Density (ρ) and thermal conductivity *(k)* of the SH PDMS + HGMs composites. **F)** Temperature profiles, including those obtained from finite element analysis (FEA), of the encapsulated battery with heater (BwH) during an overheating simulation. (inset) Photograph illustrating the simulation test setup for an overheating lithium-polymer (LiPo) battery (based on resistive heating). **G)** Temperature distributions, obtained from FEA, of the encapsulated BwH during an overheating simulation, at t = 180s. **H)** Photographs illustrating the burning—via candle flame—of the SH PDMS (top row) and the improved flame resistance of the SH PDMS + HGMs composites (middle and bottom rows). **I)** Thermogravimetric analysis curves of SH PDMS and SH PDMS + HGM composites.

Transient plane source (TPS) measurements performed using an epidermal thermal depth sensor (e-TDS) (Figure 4D) indicate the thermal conductivity of SH PDMS and SH PDMS + HGMs composites (Figure 4E).^[51]^ The thermal conductivities of the 2.5 wt% and5.0 wt% HGMs composite materials are ∼8% and ∼25% lower, respectively, than that of SH PDMS. The densities of the 2.5 wt% and 5.0 wt% HGMs composites (Figure 4E, Table S2) similarly decrease ∼14% and ∼27%, respectively, compared to SH PDMS. These decreases in thermal conductivity enhance the thermal insulation properties. Experiments to examine the relevance in the context of safety hazards associated with an overheating battery involve SH PDMS and SH PDMS + HGMs structures that fully encapsulate a LiPo battery housing that contains a resistive heater designed to simulate thermal runaway.^[6]^ Passing 250 mA through the heater causes a significant temperature increase (Figure 4F) in the battery structure. At 180 s after supply of current, the temperature of the heater of the 2.5 wt% and 5.0 wt% HGMs composites are ∼10.2°C and ∼14.5°C higher, respectively, than that of SH PDMS, indicating that the thermally insulating nature of the HGMs helps to contain and isolate the heat to the battery structure (Figure 4G). In practical applications, the thermal insulating aspects of the HGMs can be combined with other thermal protective strategies that exploit materials, layouts, and circuit designs.^[6, 52-54]^

HGMs, as a low-density filler component, migrate to the top surface during extreme cases of heating that liquify the matrix material (Figure S11). This migration creates a heat-reflective barrier that influences the thermal stability of the SH PDMS + HGMs composites.^[55-57]^ For example, flame-retardant properties begin to emerge with increasing mass fractions of HGMs, specifically in delays for combustion when the SH PDMS + HGMs composites are exposed to a flame (Figure 4H). In the case of SH PDMS, the material rapidly ignites at t ∼ 3 s. At approximately the same time point (t ∼ 3-4 s), the 2.5 wt% HGMs composite material begins to show signs of charring on the surface, but ignition does not occur until t ∼ 7 s. The 5.0 wt% HGMs composite displays even higher levels of flame-retardant properties, igniting only after 10 s of exposure to the flame and with minimal signs of charring. These results are consistent with previous studies of HGMs as fillers in other matrix materials (e.g., polyurethane, polypropylene, ethylene-vinyl-acetate, and poly(lactic acid)).^[55-59]^ The thermal stability of SH PDMS and SH PDMS + HGMs composite materials can be further characterized by the thermogravimetric analysis (TGA) (Figure 4I) and differential thermogravimetric (DTG) measurements (Figure S12). The peak degradation temperatures of the 2.5 wt% and 5.0 wt% HGMs composite materials are 543°C and 577°C, respectively, both significantly higher than that of SH PDMS (518°C).

### 2.5. An Autonomic, Self-Healing Dynamic Covalent Elastomer with Varying Functionalities for Encapsulation of a Wireless, Skin-Interfaced Mechanoacoustic Device

The SH PDMS material and its various composite forms (e.g., TC SH PDMS, SH PDMS + MC HPC, SH PDMS + HGMs, and SH PDMS + patterned nylon mesh) can be used in encapsulating structures for a wide range of skin-interfaced bioelectronic devices. As a representative example, this section focuses on a wireless sensor that mounts on the suprasternal notch to capture mechanoacoustic signatures of health-related parameters and motions of the core body.^[3-4]^ **Figure 5**A illustrates the encapsulated device in a dual-sided construct that uses SH PDMS + HGMs (2.5 wt% HGMs) and TC SH PDMS (Figure 5A, inset) on the side of the device adjacent to the embedded thin LiPo battery (Figure 1B), which poses the highest thermal safety risk. Figure 5B highlights the TC SH PDMS component of the encapsulation structure, as the device is heated from 38°C to 49°C, showing the characteristic color change as the device exceeds the activation/switching temperature of 47°C.

**Figure 5:**
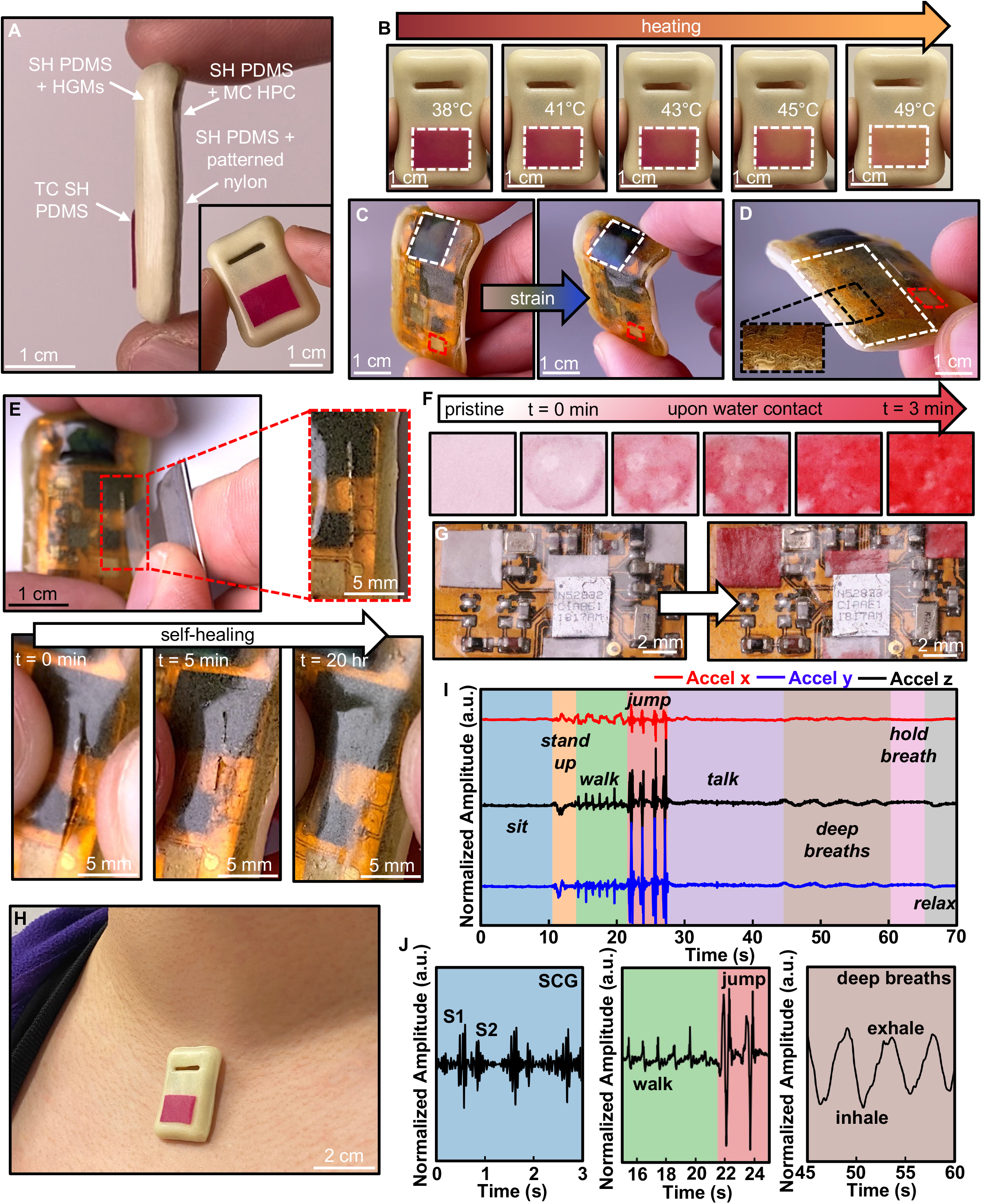
The encapsulation structure of a wireless, skin-interfaced mechanoacoustic (MA) device. **A)** Photograph depicting the side-view of a wireless, skin-interfaced MA device, encapsulated with self-healing PDMS (SH PDMS) and its variations, including thermochromic SH PDMS (TC SH PDMS), SH PDMS + hollow glass microspheres (HGMs), SH PDMS + mechanochromic hydroxypropyl cellulose (MC HPC), and SH PDMS + patterned nylon. (inset) The front view of the TC SH PDMS and SH PDMS + HGMs side. **B)** Photographs illustrating the color change of the TC SH PDMS, outlined in white, associated with heating. **C)** Photographs illustrating the color change of the SH PDMS + MC HPC, outlined in white, associated with strain from bending of the device. Outlined in red is the water contact indicator, as viewed through transparent SH PDMS. **D)** Photograph illustrating the SH PDMS + patterned nylon section, outlined in white, of the encapsulation structure. Outlined in red is the water contact indicator, as viewed through transparent SH PDMS. **E**) Photographs illustrating the self-healing behavior of the encapsulation structure. **F)** Time-lapse of the color changes associated with water exposure to the water contact indicator material. **G)** Photographs of the water contact indicators overlaid on top of the printed circuit board, changing color as water is introduced. **H**) Photograph of the encapsulated MA device on the suprasternal notch of an adult female subject. **I)** Representative triaxial accelerometry signals obtained from the encapsulated MA device, worn by an adult female subject performing various physical activities. **J)** Z-axis accelerometry signals illustrate unique signatures of different body processes, including: cardiac activity, in the form of the seismocardiogram (SCG) waveform with distinct S1 and S2 heart sounds (left), physical motions such as walking and jumping (middle), and respiration waveforms during deep breaths (right).

The other side uses SH PDMS + MC HPC, SH PDMS + patterned nylon, and SH PDMS. The SH PDMS + MC HPC resides at the location of the device (Figure 5C) that experiences the largest mechanical deformations, typically bending associated with removal of the device from the skin. The remaining area uses SH PDMS + patterned nylon (Figure 5D, outlined in white) to increase mechanical robustness, with small windows of transparent SH PDMS for purposes described below (Figures 5C-D, outlined in red). The encapsulation structure demonstrates autonomic self-healing behavior (Figure 5E), in response to a severe cut that exposes the internal electronic components. A significant portion of the cut self-heals after only 5 mins, with full healing after 20 h.

In addition to direct exposure of these components to the skin, an additional safety hazard follows from electrical shock mediated by water or biofluids. Thin hydrochromic paper materials inserted into the encapsulating package at locations visually observable through transparent windows can serve as useful indicators of water exposure. In the example reported here, as water reaches a commercial indicator of this type, its color changes from white to red (Figures 5F-G).

The device mounted on a volunteer subject appears in Figure 5H. Triaxial accelerometry (Figure 5I) and gyroscopic measurements (Figure S13) at this location allow for continuous capture of various signals including those associated with cardiac cycles (represented in the form of seismocardiograms (SCGs)), large physical motions (e.g., walking and jumping), and respiratory activity (both inhalation and exhalation) derived from the acceleration data in the direction normal to the surface of the skin (Figure 5J). The SCG waveforms clearly exhibit the characteristic S1 and S2 heart sounds, indicating that the encapsulating structure of the device does not interfere with or dampen such sensitive physiological data collection (Figure S14).

## 3. Conclusion

The collective results presented here suggest that advanced materials strategies can be used in wireless, skin-interfaced bioelectronic devices to manage safety risks in a manner that can complement and extend capabilities based on traditional sensor and electronic circuit methods. Specifically, a self-healing formulation of a silicone polymer serves as a matrix material in soft encapsulating structures that offer various stimuli-responsive properties, such as reversible thermo/mechanochromism and strain-adaptive stiffening, in addition to other improved thermal-related characteristics. This multifunctional material construct addresses various risks in mechanical, thermal, and electrical modes of device failure. Demonstrations of use with a mechanoacoustic sensor platform suggest wide-ranging applicability across other types of body-integrated bioelectronic systems. Extending these ideas to other levels of materials function in safety and optimizing these strategies for cost-effective manufacturing represent promising directions for future research.

## 4. Experimental Section/Methods

### Synthesis of Self-Healing Poly(dimethylsiloxane) (SH PDMS)

Starting materials and all commercially available solvents and reagents were used as received, without further purification. First, 1,3,5-triformylbenzene (TFB) (Thermo Scientific) was dissolved in N-N-dimethylformamide (DMF) (Sigma-Aldrich) to form a 0.33 M solution. This solution was then vortex mixed with α,ω-telechelic aminopropyl PDMS (M_w_ ∼ 5000 g mol^-1^, DMS-A21, Gelest) at a 1:1 molar ratio in a 20 mL glass scintillation vial for 3 min. The resulting mixture was poured into a Teflon dish and placed into a vacuum oven at 50°C for 24 h for crosslinking and drying, which eventually yielded a yellow, transparent, and self-healing elastomer (SH PDMS).

### Synthesis of Thermochromic SH PDMS (TC SH PDMS)

Starting materials and all commercially available solvents and reagents were used as received, without further purification. First, pre-synthesized SH PDMS films were re-processed by dissolving them in chloroform (Sigma-Aldrich). After dissolution, a magenta thermochromic (TC) leuco dye powder (LCR Hallcrest), with an activation/switching temperature at 47°C, was added to the solution at varying mass fractions (0.1-0.5 wt%). The dye was pulverized by a mortar and pestle before use. The solution with the dye powder was then thoroughly mixed by using a vortex mixer. The mixture was poured into a target well to control the film thickness. Overnight drying at room temperature yielded a magenta TC SH PDMS elastomer.

### Characterization of TC SH PDMS

Thin (∼300 μm) films of TC SH PDMS, with varying mass fractions of TC leuco dye, were placed on a hot plate and heated to 55°C. Videos were captured of the samples during, and the changes in color were analyzed via ImageJ software to determine the changes in RGB values, as well as the response times.

### Synthesis of Mechanochromic (MC) Liquid Crystalline Hydroxypropyl Cellulose (HPC)

Starting materials and all commercially available solvents and reagents were used as received, without further purification. A 60 wt% aqueous solution of hydroxypropyl cellulose (HPC) (M_w_ ∼ 100,000 g mol^-1^, Sigma-Aldrich), with 0.5 wt% of black silicone dye (Silc Pig, Smooth-On), was formed by mixing in a planetary centrifugal mixer (Thinky ARE-310) for 30 min at 2000 RPM. The resulting mixture was allowed to rest in a closed container placed inside a humid environment to prevent drying. This process yielded an iridescent, amber-hued and mechanochromic (MC) HPC gel.

### Integration of MC HPC with SH PDMS

The synthesized MC HPC gel was cast onto the middle region of a thin (∼50 μm) SH PDMS layer via a piping bag with a 1-mm diameter opening. Another thin (∼50 μm) SH PDMS layer was carefully placed on top of the deposited MC HPC gel and the bare edges of the top and bottom SH PDMS films were pressed against each other and permanently bonded via autonomic self-healing and self-adhesion. This process produced a fully sealed and encapsulated MC HPC gel within the SH PDMS layers.

### Synthesis of SH PDMS + Hollow Glass Microsphere (HGM) Composites

Starting materials and all commercially available solvents and reagents were used as received, without further purification. First, 1,3,5-triformylbenzene (TFB) (Thermo Scientific) was dissolved in N,N-dimethylformamide (DMF) (Sigma-Aldrich) to form a 0.33 M solution. This solution was then vortex mixed with α,ω-telechelic aminopropyl PDMS (M_w_ ∼ 5000 g mol^-1^, DMS-A21, Gelest) at a 1:1 molar ratio, with varying mass fractions (0.25-0.50 wt%) of hollow glass microspheres (HGMs) (Q-Cel 300, Potters Industries), in a 20 mL glass scintillation vial for 3 min. The resulting mixture was poured into a Teflon dish and placed into a vacuum oven at 50°C for 24 h for crosslinking and drying. The result was a whitish, self-healing elastomer composite (SH PDMS + HGMs).

### Fourier Transform Infrared (FTIR) Spectroscopy

Solid state FTIR transmission spectra were measured with the use of a Thermo Scientific Nicolet iS50, over a range of 4000 to 400 cm^-1^. FTIR of SH PDMS (cm^-1^): ν = 2962 (w; ν(C–H)), 1650 (w; ν(C=N)), 1257 (m; ν(C–H)), 1009 (s; ν(Si–O–Si)), 786 (s; ν(Si–C)).

### Scanning Electron Microscopy (SEM)

Samples were first coated with 18 nm of osmium using a Filgen Osmium Plasma Coater. Surface morphologies of coated samples were then characterized with a Hitachi SU-8030 SEM, with an acceleration voltage of 5.0 kV and a working distance of 12.5 mm.

### UV-Vis Spectroscopy

Solid state UV-Vis absorption and transmission spectra were measured with a Perkin Elmer LAMBDA 1050 spectrophotometer, over a range from 200 to 800 nm.

### Design and Characterization of Mechanically-Reinforcing and Strain-Adaptive Stiffening Meshes

Poly-L-lactic acid (PLLA, #233-146-54), poly(methyl methacrylate) (PMMA, #676-315-97), cellulose acetate (CA, #502-330-14), and nylon-6 (#142-214-93) films were purchased from Goodfellow Corporation. Poly(ethylene terephthalate) (PET, #P04DW0912) was purchased from Grafix Plastics. Details for the basic design principles of the patterned mesh networks for mechanical reinforcement and strain-adaptive stiffening of SH PDMS can be found elsewhere.^[39]^ Desired reinforcing patterns were defined by a laser cutter system (ProtoLaser R, LPKF). Stress/strain profiles were determined from uniaxial tensile tests (strain rate = 0.5 mm s^-1^) with a dynamic mechanical analyzer (RSA-G2 Solids Analyzer, TA Instruments).

### FEA of Mechanically-Reinforcing and Strain-Adaptive Stiffening Meshes

3D FEA performed in the commercial software package Abaqus captured the nonlinear deformations of the mechanically-reinforcing and strain-adaptive stiffening meshes. Experimentally measured mechanical properties of the constitutive material (i.e., nylon-6) were adopted in the analyses. Four-node shell elements were used to simulate the mesh-type structures, with refined finite element meshes for accuracy. Linear buckling analyses for the mesh structures determined the critical buckling strains and corresponding buckling modes, which were then implemented as initial geometric imperfections in post-buckling simulations. Deformed configurations and strain distributions for the mesh structures were thus obtained.

### Preparation of Silk/Glycerol Blend Films

An aqueous silk solution was formed using *Bombyx mori* silkworm cocoon following the previous study.^[60]^ After the degumming and dissolution in 9.3 M of LiBr solution, the solution was dialyzed in deionized (DI) water with a dialysis flask (Thermo Scientific) for 7 days. The final concentration was 6.5% (w/v). The solution was mixed with glycerol (Sigma-Aldrich) for plasticization. Glycerol was added to the solution at weight ratios of 0%, 10%, 20%, 30%, 40%, and 50% and mixed. The mixed solution was poured into a petri dish and dried overnight at room temperature to obtain the glycerol-plasticized silk (Silk/Gly) films.^[61]^

### Integration of Patterned Serpentine Nylon Mesh with SH PDMS

The patterned serpentine nylon mesh was placed on top of a layer of SH PDMS (thickness ∼500 μm). Another SH PDMS layer (thickness ∼500 μm) was carefully placed on top of the nylon mesh/SH PDMS and the exposed surfaces of the top and bottom SH PDMS films were pressed together and permanently bonded via autonomic self-healing and self-adhesion. The result was a patterned nylon mesh fully integrated within the SH PDMS layers.

### Mechanical Characterization of SH PDMS, SH PDMS + HGM Composites, and SH PDMS + Patterned Nylon

Stress/strain profiles were determined from uniaxial tensile tests (strain rate = 0.1 mm s^-1^) with a dynamic mechanical analyzer (RSA-G2 Solids Analyzer, TA Instruments).

### Digital Image Correlation (DIC) for Self-Healing Analysis

The experiments involved recording a set of uniaxially-stretched SH PDMS films using a high-speed camera (2048 × 1088 resolution, HT-2000M, Emergent) with 35 mm imaging lenses (F1.4 manual focus, Kowa) at the frame rate of 100 fps. A slit film of SH PDMS was investigated at two different healing times: 1) immediately after attaching the slit portions together, and 2) after 15 mins of self-healing. The film was uniformly coated with black speckles (Figure S1) by the spray-painting method. The opensource 2D-DIC software, Ncorr, was used to measure material deformation of the self-healed film.^[62]^ To achieve high-resolution and accurate deformation characteristics, the DIC subset radius and spacing were set as 10 pixels and 5 pixels, respectively, resolving over 1500 displacement grids with the grid resolution of ∼400 μm. The strain magnitude and shear strain were computed based on the Triangular Cosserat Point Theory.^[63]^

### Digital Image Correlation (DIC) for MC HPC Analysis

Fluidic channel molds with a serpentine pattern (inner radius = 3.5 mm, cross-sectional width = 3.0 mm, height = 2.0 mm, length = 65.0 mm) were prepared by 3D-printing (printer: Form 3, Formlabs, material: Clear V4). These molds were used for full encapsulation of MC HPC gel within the elastomer channels. Resulting color changes of the MC HPC system, induced by uniaxial stretching, were compared with strain values as measured by DIC.

### FEA for MC HPC Analysis

The commercial FEA software ABAQUS was used to determine the mechanical performance of SH PDMS + MC HPC when subjected to uniaxial strain. A 3D rectangular model with dimensions of 70 mm x 30 mm x 6 mm was used for the fluidic channel mold, which includes the serpentine pattern of the MC HPC with a thickness of ∼2 mm. Displacement boundary conditions were imposed at the ends of the 3D rectangular elastomer encapsulation model to stretch uniaxially along the longitudinal direction by up to 30%. The strain ε and in-plane displacement U_x_ and U_y_ fields were computed at 30% uniaxial stretch (Figures S5-S6), where the simulation closely matches the DIC field measurements. The elastomer encapsulation and MC HPC were modeled by hexahedron elements (C3D8R) and the total number of elements used in the simulation is 125,000.

### Thermal Conductivity Measurements of SH PDMS and SH PDMS + HGM Composites

A thin epidermal thermal depth sensor (e-TDS) (R = 1.5 mm, q = 2 mW mm^-2^) placed onto the surface of the test sample enabled measurements of the thermal conductivity (*k*), via the transient plane source (TPS) technique (heating time (t) = 30 s).^[51]^ Presented results are the average and standard deviation of 3 consecutive measurements per sample.

### Density Measurements of SH PDMS and SH PDMS + HGM Composites

The density of the material of interest was determined by dividing the mass by the cubic volume of rectangular film samples. The density of the filler HGMs was provided by the manufacturer (Potters Industries). To determine the theoretical densities of the SH PDMS/HGM composite materials, the volume fraction of the filler was first calculated using the known mass fractions and densities of the filler and matrix:

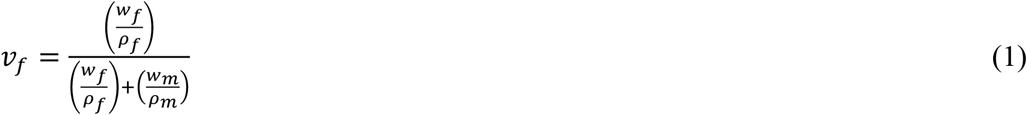

where *ν*_*f*_, *w*_*f*_,*ρ*_*f*,_ *w*_*m*_,*ρ*_*m*_ represent the filler volume fraction, filler mass fraction, filler density matrix mass fraction, and matrix density, respectively.

Then, the rule of mixtures was used to determine the theoretical density of the composite material:

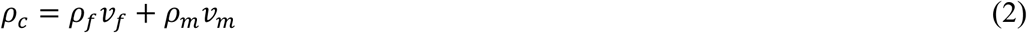

where *ρ*_*c*_,*ρ*_*f*_, *ν*_*f*,_*ρ*_*m*,_*ν*_*m*_ represent the composite material density, filler density, filler volume fraction, matrix density, and matrix volume fraction. Presented results are the average and standard deviation of 3 measurements taken per sample.

### Overheating Battery Simulation

The process for simulating an overheating battery (thermal runaway) appears elsewhere.^[6]^ Briefly, 250 mA of current was applied to a SH PDMS or SH PDMS + HGM composite-encapsulated battery enclosure with heater (BwH) device, consisting of a lithium polymer (LiPo) battery enclosure (DNK201515, DNK Power) with a copper resistive heating element. The resulting temperature increase inside the BwH was monitored via measurement of the voltage drop across the BwH.

### FEA of overheating battery

The commercial software COMSOL Multiphysics revealed aspects of heat conduction and convection from the BwH and the encapsulation material. The heat generated by Joule heating conducted through the battery and the encapsulation material. The BwH and encapsulation material experienced natural convection of air at room temperature (T_air_ = 22°C), which were introduced as thermal-fluid boundary conditions in the FEA model. To compare thermal insulation functions of different encapsulation materials, the heat generation from the heater was kept constant by setting the current as 250 mA. Higher heater temperature indicates that more heat is trapped in the battery by the encapsulation material with a higher thermal impedance (R_encapsulation material_ = (T_heater_ – T_air_)/Q – R_nc_). FEA results of heater temperatures are summarized in Figures 4F-G.

Thermophysical properties of the BwH were analyzed from the heater and surface temperatures as a function of the heater power without the encapsulation material. The effective heat capacity and the density are average properties weighted by the mass of electrodes and electrolytes. Effective thermophysical properties of the SH PDMS + HGM composite materials were obtained from experiments using the TPS method, as previously described.

To verify the mesh independence and time step independence, different models with mesh densities ranging from 12 elements per cubic millimeter to 20 elements per cubic millimeter and time steps ranging from 0.05 s to 0.1 s were analyzed. A temperature difference of < 0.1°C between these models confirmed that the mesh density of 12.3 elements per cubic millimeter and time step of 0.1 s were selected for the FEA analysis. Additionally, effective heat capacities of the SH PDMS + HGM composite materials were estimated by comparing FEA results with experimental results.

### Thermogravimetric Analysis

Thermal stability of SH PDMS and SH PDMS + HGM composite materials was determined with a thermogravimetric analyzer (STA 449, Netzsch) over a temperature range from 30°C to 700°C, at a heating rate of 20°C min^-1^ under nitrogen atmosphere.

### Device Encapsulation

A wireless, skin-interfaced mechanoacoustic sensor, as previously reported^[3-4]^, was encapsulated with the SH PDMS elastomer and various composites (e.g., TC SH PDMS, SH PDMS + HGMs, SH PDMS + MC HPC, SH PDMS + patterned nylon), with the use of concave and convex aluminum molds. Layers of SH PDMS and associated composites were permanently bonded to each other via autonomic self-healing and self-adhesion, which eventually yielded a fully encapsulated device within the SH PDMS-based materials.

### Device Application, Data Collection, and Data Processing

The encapsulated mechanoacoustic device was applied onto the suprasternal notch of a 27-year-old female subject and secured with a medical-grade, double-sided silicone/acrylate adhesive (2477P, 3M). The device was wirelessly connected to an iPhone with a customized application for data transmission, collection, and storage. Triaxial accelerometry and gyroscopic measurements were downloaded from the iPhone’s internal memory. Z-axis accelerometry data were bandpass filtered (15-30 Hz) to extract seismocardiogram (SCG) waveforms.

## Acknowledgements

C.L., J.-T.K., D.S.Y., and D. C. contributed equally to this work. C.L., S.R.M., and R.A. acknowledge funding support from the National Science Foundation Graduate Research Fellowship Program (NSF DGE-1842165). R.A. acknowledges funding support from the Ford Foundation Predoctoral Fellowship. This work made use of the Keck-II facility and the NUFAB facility of Northwestern University’s NUANCE Center, which has received support from the SHyNE Resource (NSF ECCS-2025633), the IIN, and Northwestern’s MRSEC program (NSF DMR-1720139). Engineering efforts were supported by the Querrey Simpson Institute for Bioelectronics at Northwestern University. The authors gratefully acknowledge Seojin Yoo for her contributions to this work.

