## Supporting Information for "Multifunctional Materials Strategies for Enhanced Safety of Wireless, Skin-Interfaced Bioelectronic Devices"

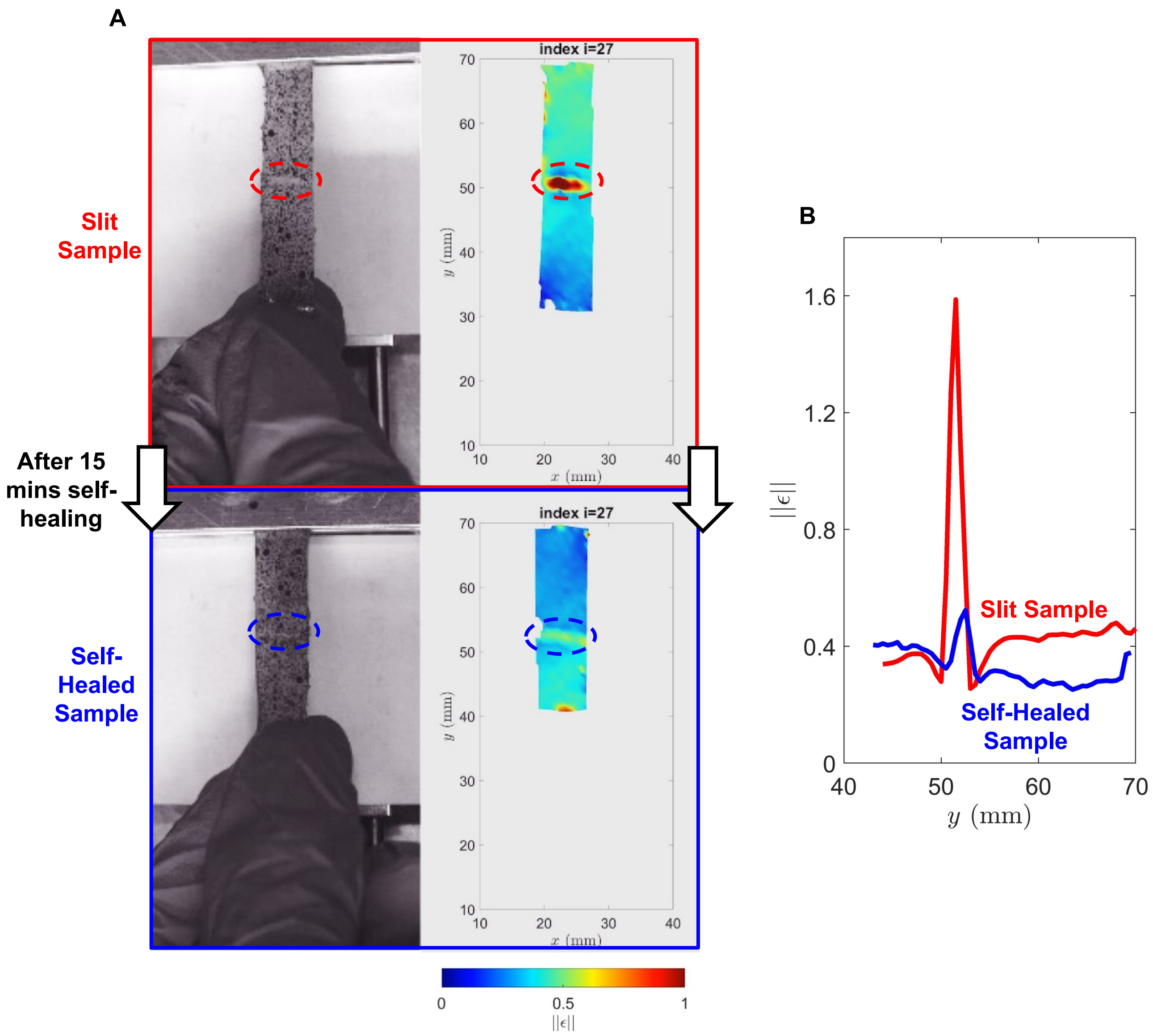


**Figure S1: Digital image correlation (DIC) analysis of the autonomic self-healing behavior of the dynamic covalent elastomer (SH PDMS). A)** Photographs (left column) and the corresponding strain distribution maps (right column)—via DIC experiments—of a slit film of SH PDMS undergoing uniaxial strain (top row, red), which is then allowed to self-heal for 15 mins (bottom row, blue) and undergo uniaxial strain again. **B)** Measured strain magnitude at the slit (red) and self-healed (blue) region of the SH PDMS film.

| Material | Efficiency after 15 min (%) | Efficiency after 24 h (%) |
|---|---|---|
| SH PDMS | 32.2 | 99.9 |
| TC SH PDMS | 49.7 | 92.2 |
| 2.5 wt% HGMs composite | 6.3 | 98.9 |
| 5.0 wt% HGMs composite | 26.0 | 82.0 |

**Table S1. Self-Healing Efficiencies "η" of SH PDMS and SH PDMS composite materials at different time points.**

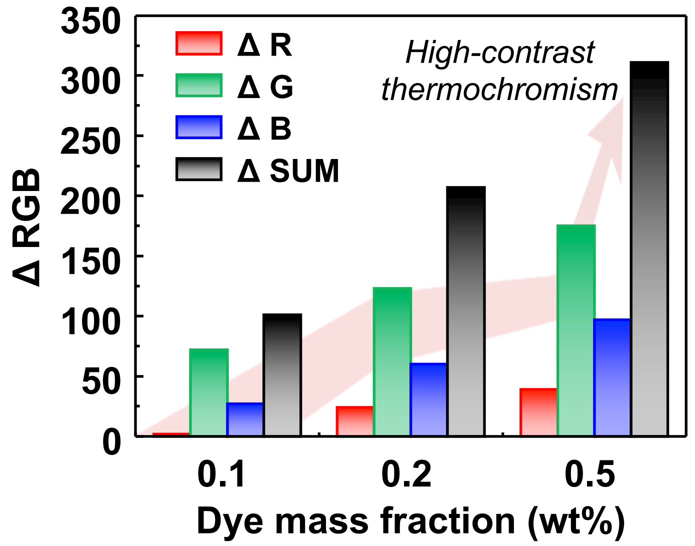


**Figure S2: Thermochromic properties, as illustrated by the change in RGB values between the relaxed and heated states of the dynamic covalent elastomer matrix integrated with leuco dye (TC SH PDMS), with varying mass fractions.**

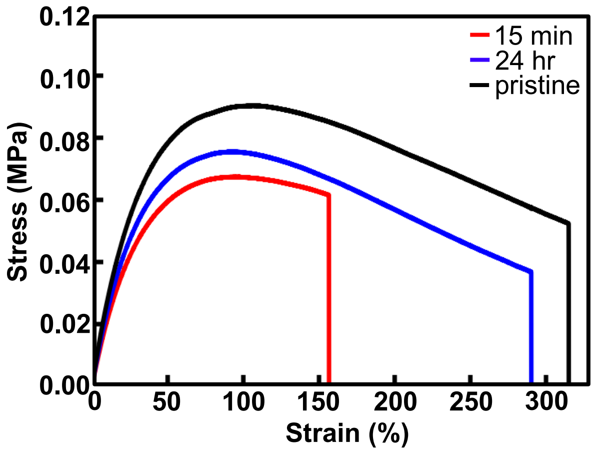


**Figure S3: Stress/strain curves of the dynamic covalent elastomer matrix integrated with thermochromic leuco dye (TC SH PDMS), including those after different time periods of self-healing.**

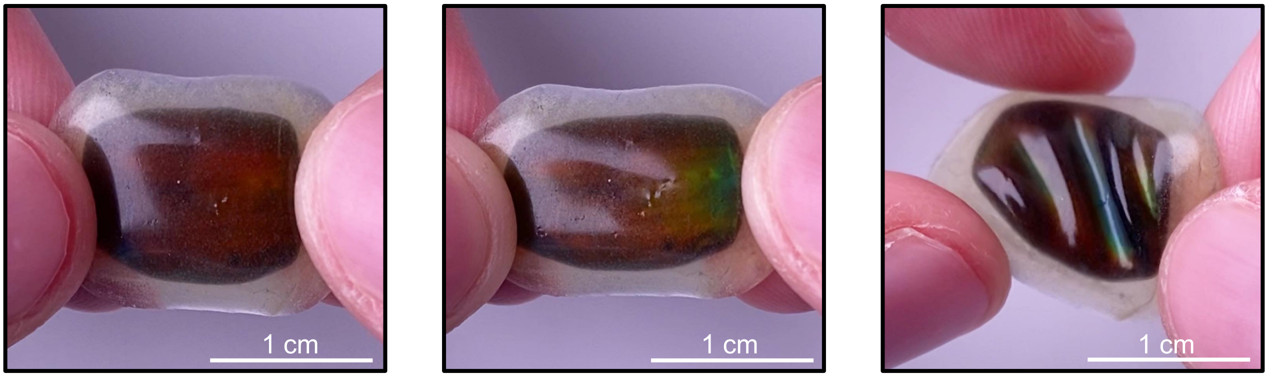


**Figure S4: The dynamic covalent elastomer (SH PDMS) with integrated mechanochromic hydroxypropyl cellulose (MC HPC).** Photographs illustrating the integration of MC HPC within SH PDMS elastomer layers. The initial state of the SH PDMS + MC HPC composite material exhibits an amber/red color (left). As uniaxial strain is slowly applied (middle), a blue-shift occurs due to the pitch decrease of the cholesteric liquid crystalline structure of HPC and green/blue colors are observed in the material. Applied pressure/stress from bending/folding of the material (right) also induces a blue-shift, as green/blue colors are observed along the folded/bent regions of the material.

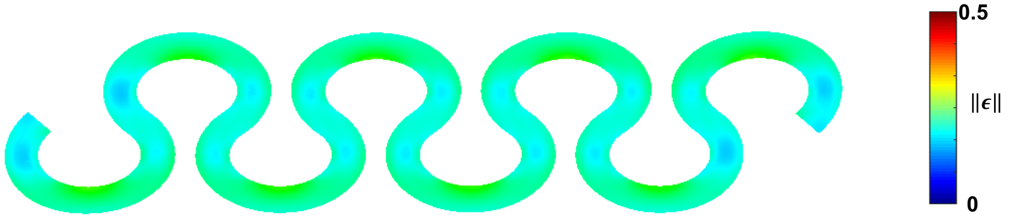


**Figure S5: Finite element analysis (FEA) of the strain distribution field of the dynamic covalent elastomer integrated with mechanochromic hydroxypropyl cellulose (SH PDMS + MC HPC).**

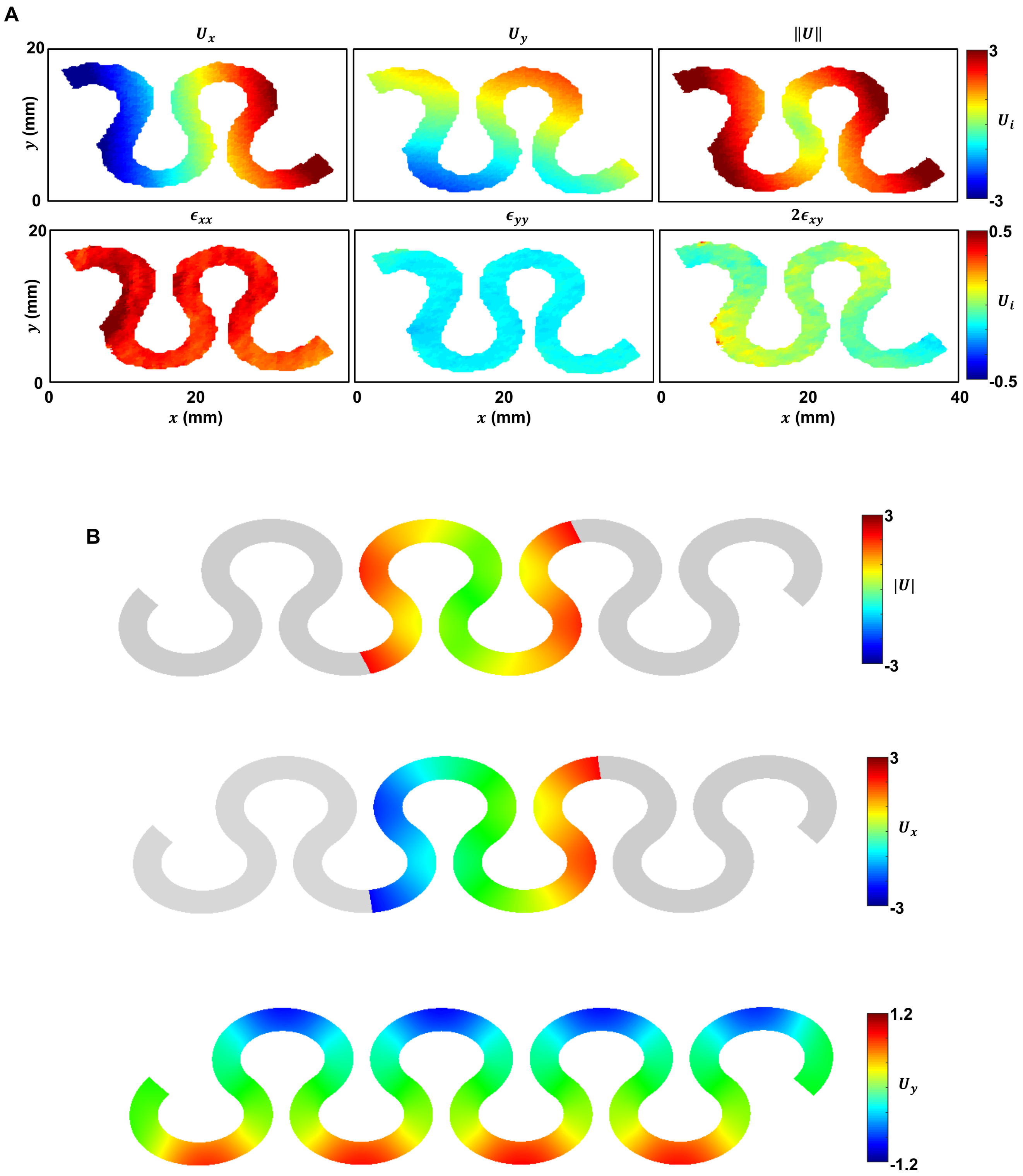


**Figure S6: Displacement fields of the uniaxially-strained dynamic covalent elastomer integrated with mechanochromic hydroxypropyl cellulose (SH PDMS + MC HPC) by: A) digital image correlation (2D-DIC), and B) finite element analysis (FEA).**

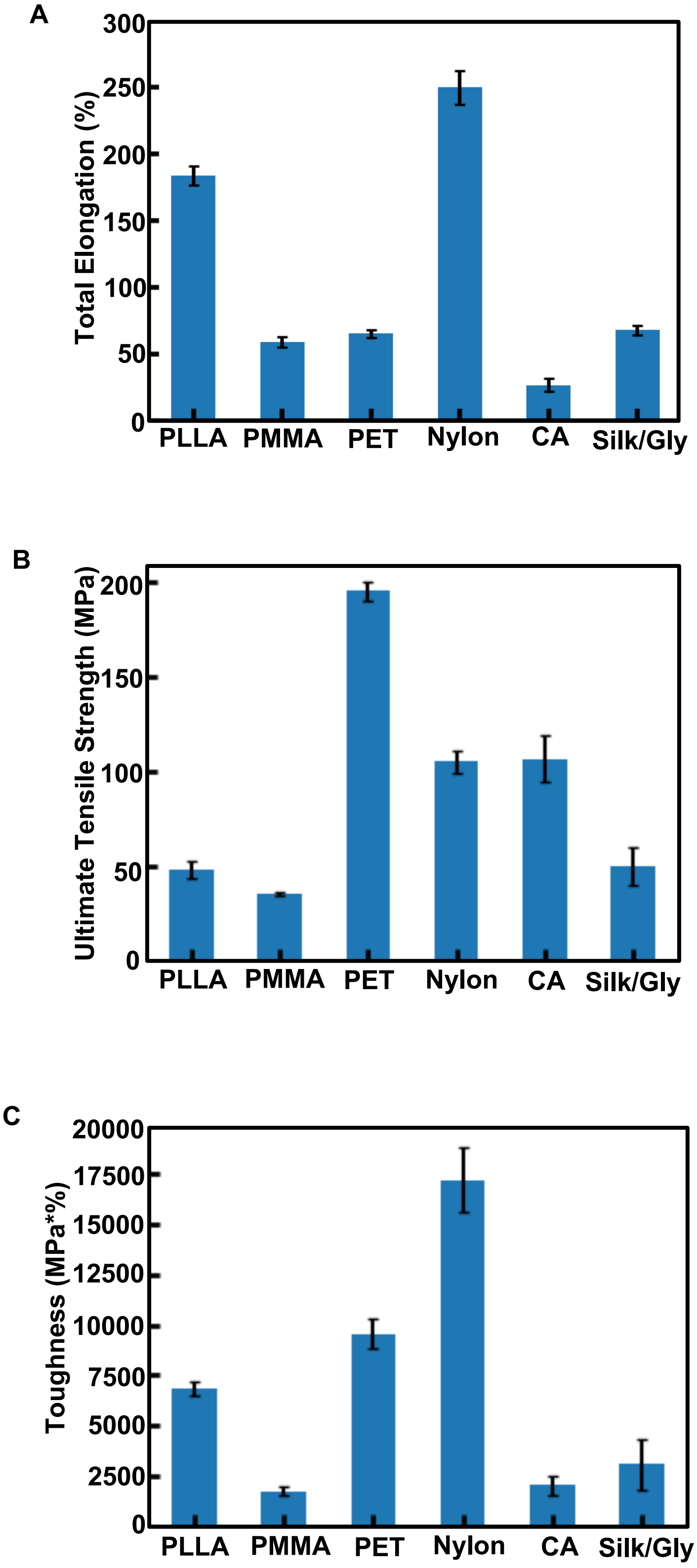


**Figure S7: Measurements from dynamic mechanical analysis of A) total elongation, B) ultimate tensile strength, and C) toughness, of various commercially available or natural polymers.**

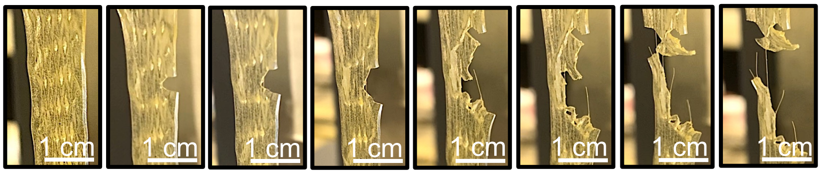


**Figure S8: Photographs illustrating the tear/fracture behavior of the SH PDMS + patterned nylon mesh composite material.**

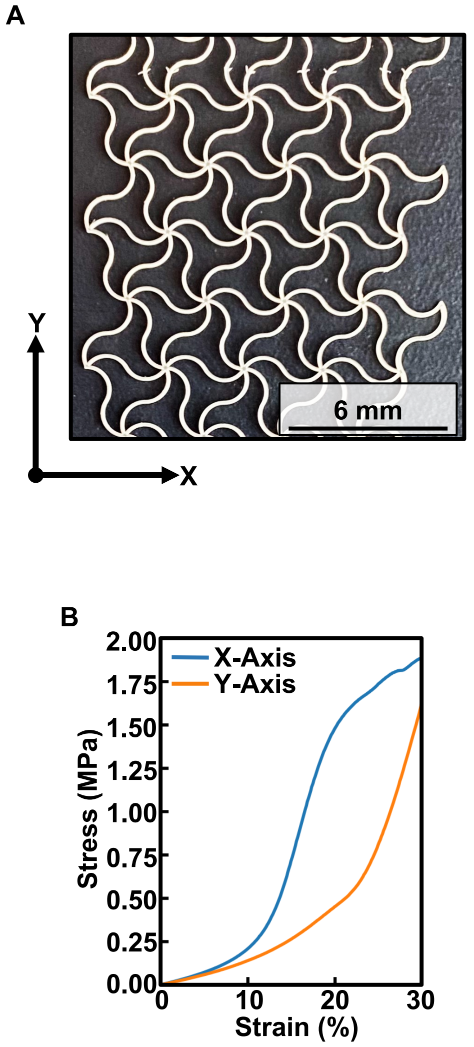


**Figure S9: Anisotropic mechanical behavior of the (A) patterned nylon mesh in a triangular serpentine layout, illustrating (B) different strain threshold limits when subject to strain along the X-Axis vs. the Y-Axis.**

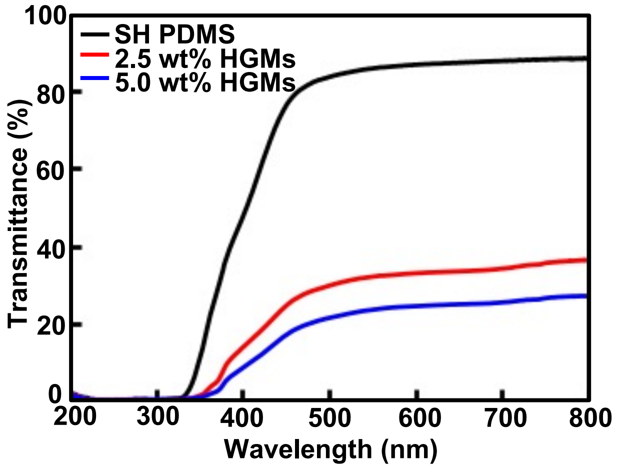


**Figure S10: UV-Vis transmittance spectra of the dynamic covalent elastomer (SH PDMS) and the dynamic covalent elastomer integrated with varying mass fractions of hollow glass microspheres (HGMs) (SH PDMS + HGMs).**

| Material | Measured Density (g cm$^{-3}$) | Theoretical Density (g cm$^{-3}$) |
|---|---|---|
| SH PDMS (0 wt% HGMs) | 0.947 | n/a |
| HGMs | 0.120 | n/a |
| 2.5 wt% HGMs composite | 0.819 | 0.808 |
| 5.0 wt% HGMs composite | 0.689 | 0.704 |

**Table S2. Comparison of measured and theoretical densities of the dynamic covalent elastomer (SH PDMS) and hollow glass microsphere (HGM) composite materials.**

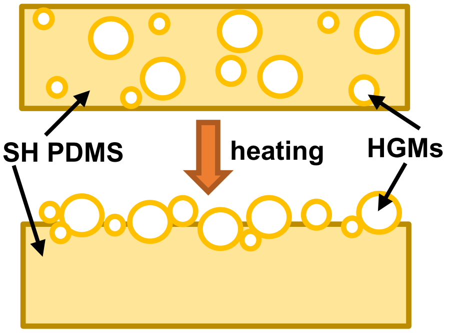


**Figure S11. Schematic diagram illustrating the HGM migration process upon heating.**

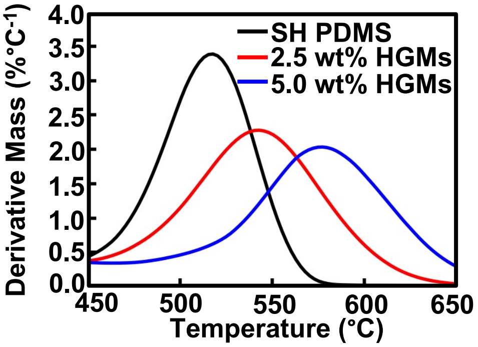


**Figure S12: Differential thermogravimetric (DTG) curves of SH PDMS and SH PDMS + HGMs composite materials.**

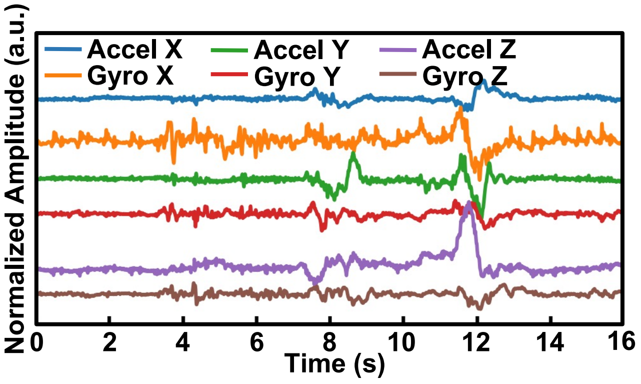


**Figure S13: Representative triaxial accelerometry and gyroscope signals as obtained from the encapsulated mechanoacoustic device, worn by an adult female subject.**

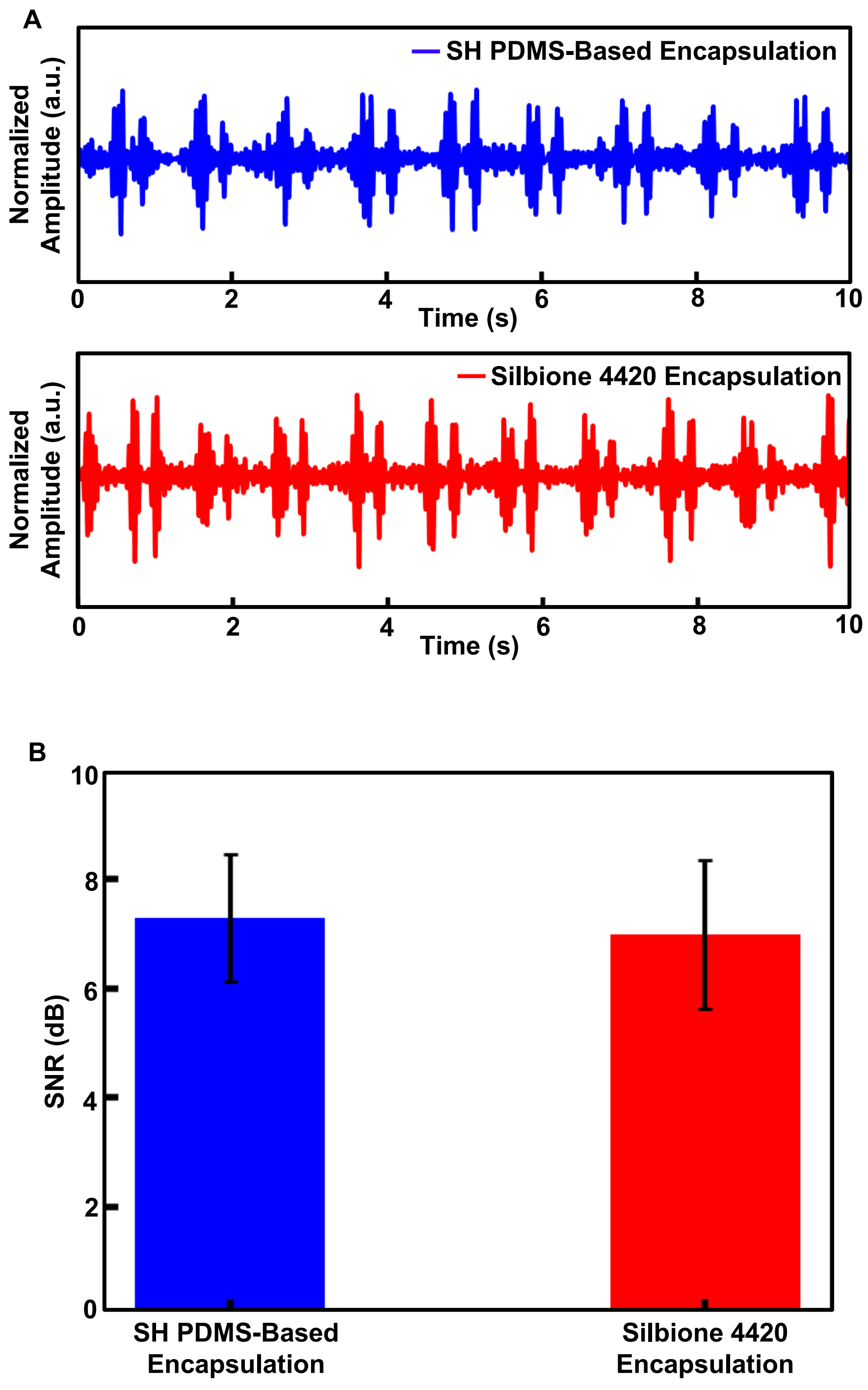


**Figure S14: A) Representative seismocardiograms (SCGs) derived from Z-axis accelerometry signals as obtained from the mechanoacoustic (MA) device encapsulated with varying materials, including SH PDMS-based materials and a conventional medical-grade silicone, Silbione 4420. B) Signal-to-noise ratio (SNR) of obtained accelerometry data from the MA device encapsulated with varying materials, , including SH PDMS-based materials and a conventional medical-grade silicone, Silbione 4420.**
